# A specific phosphorylation-dependent conformational switch of SARS-CoV-2 nucleoprotein inhibits RNA binding

**DOI:** 10.1101/2024.02.22.579423

**Authors:** Maiia Botova, Aldo R. Camacho-Zarco, Jacqueline Tognetti, Luiza Mamigonian Bessa, Serafima Guseva, Emmi Mikkola, Nicola Salvi, Damien Maurin, Torsten Herrmann, Martin Blackledge

**Affiliations:** Univ. Grenoble Alpes, CNRS, CEA, IBS, F-38000 Grenoble; Department of Biochemistry and Molecular Biophysics, Columbia University, New York, USA; Evotec (France) SAS, Toulouse, France; Sanofi R&D, Integrated Drug Discovery, 94403 Vitry-sur-Seine, France

## Abstract

The nucleoprotein (N) of SARS-CoV-2 encapsidates the viral genome and is essential for viral function. The central disordered domain comprises a serine-arginine-rich domain (SR) that is hyperphosphorylated in infected cells. This modification is thought to regulate function of N, although mechanistic details remain unknown. We use time-resolved NMR to follow local and long-range structural changes occurring during hyperphosphorylation by the kinases SRPK1/GSK-3/CK1, thereby identifying a conformational switch that abolishes interaction with RNA. When 8 approximately uniformly-distributed sites are phosphorylated, the SR domain competitively binds the same interface as single-stranded RNA, resulting in RNA binding inhibition. Phosphorylation by PKA does not prevent RNA binding, indicating that the pattern resulting from the physiologically-relevant kinases is specific for inhibition. Long-range contacts between the RNA-binding, linker and dimerization domains are also abrogated, phenomena possibly related to genome packaging and unpackaging. This study provides insight into recruitment of specific host kinases to regulate viral function.

## Introduction

Severe acute respiratory syndrome coronavirus 2 (SARS-CoV-2), the infectious agent responsible for the recent Covid-19 pandemic, is an enveloped positive-sense single-stranded RNA virus of the betacoronavirus genus that expresses its own replication machinery. Genome replication is achieved by the RNA-dependent RNA polymerase, whose structure has been investigated by cryo-electron microscopy (*1–3*), within viral replication organelles called double membrane vesicles (DMVs) that are remodelled from the endoplasmic reticulum (*4*). DMVs have been shown to constitute principal sites of viral RNA replication (*5*, *6*), containing narrow exit channels formed by non-structural viral proteins (nsps) through which newly synthesized single stranded RNA can exit prior to encapsidation.

The nucleoprotein is the most abundant protein in beta-coronaviruses, and an important cofactor of the replication machinery (*7*, *8*) that is known to colocalize to the replication transcription complex (*9–11*) via an essential interaction with the N-terminus of non-structural protein 3 (nsp3) (*12*, *13*). SARS-CoV-2 nucleoprotein (N) binds to, and encapsidates, the viral genome, shielding the RNA from the host innate immune system, as well as playing an essential but still poorly understood role in regulation of gene transcription (*14*). Electron tomography was used to identify N-RNA complexes within SARS-CoV-2 virions, displaying a morphology resembling beads on a string (*15*, *16*).

N is highly disordered as revealed by NMR spectroscopy (*17*, *18*), comprising three intrinsically disordered domains (N1, N3 and N5), flanking the RNA-binding domain (N2) and the dimerization domain N4 (figure 1), whose structures have both been determined at atomic resolution (*19–23*). RNA binding to N2 has been described (*24–31*), including its potential role in liquid-liquid phase separation of N-RNA mixtures (*26*, *27*, *32–38*). N4 forms a highly stable dimeric structure involving two domain-swapped β-hairpins. Although N2 is known to represent the main RNA binding domain, secondary RNA binding sites have been proposed to exist in both N3 (*39*, *40*, *26*, *36*) and N4 (*39*, *41–45*), an observation supported by assembly of truncation mutants of N into large supramolecular complexes of the dimensions of RNPs (*46*). N5 has also been associated with assembly into higher order oligomeric states (*47*).

**Figure 1.**
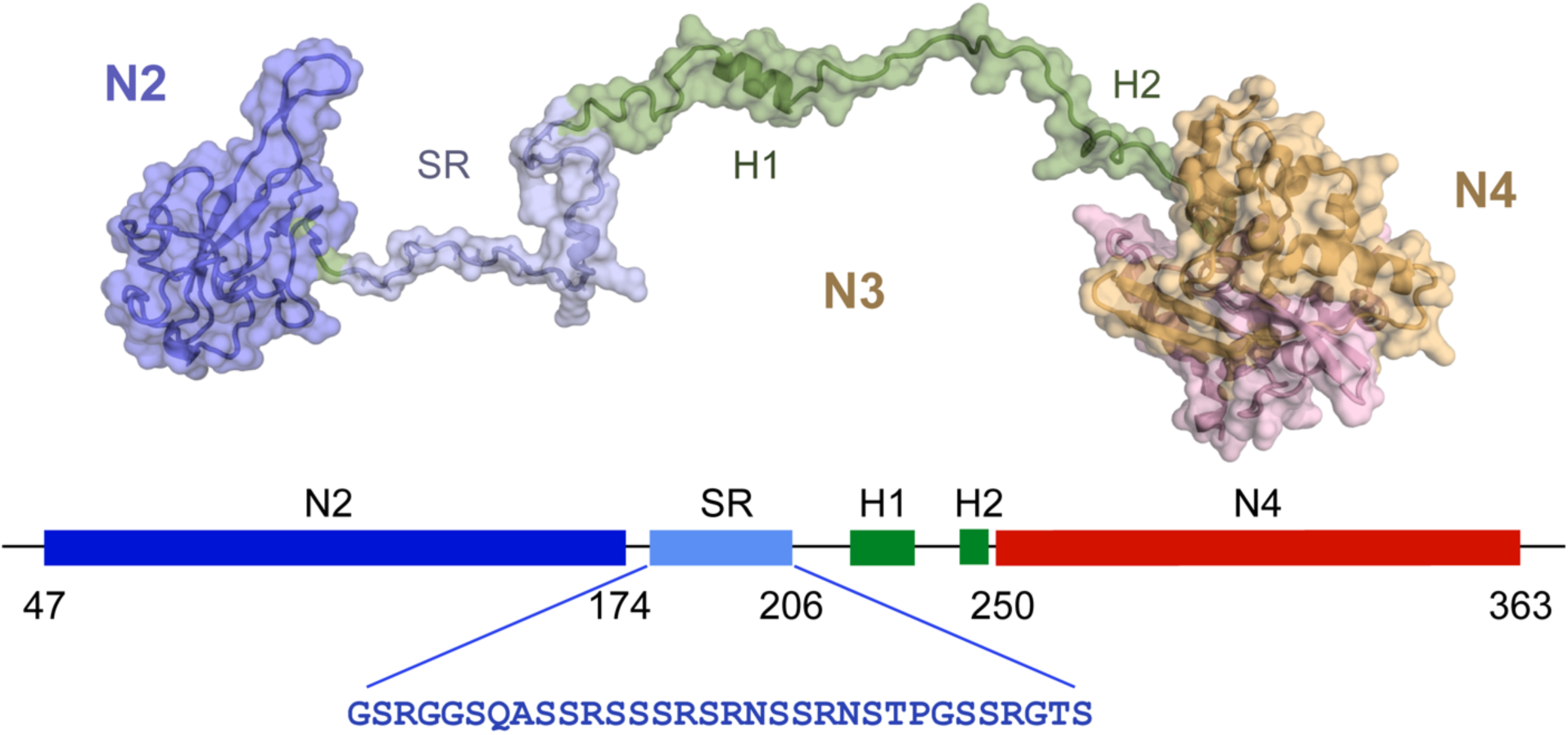
Figurative representation of SARS-CoV-2 nucleoprotein (N). SARS-CoV-2 nucleoprotein comprises 5 domains (N1-N5). The construct shown here comprises three of these domains, N2, the RNA binding domain, N3, the disordered central domain containing two helices and the Serine-Arginine-rich SR region that is hyperphosphorylated in infected cells, and the dimerization domain N4. The sequence of the SR region is shown in blue.

N3 is of particular interest for a number of reasons. It comprises three distinct linear segments, a serine/arginine rich region (SR region, S176-S206), a hydrophobic, Leucine-rich helix (H1) and a polar region, comprising a glutamine-rich strand and terminated by a Lysine-rich region that forms a detached helix (H2) in the crystal structure of N4, and represents the junction between these two domains. N3 was recently shown to fold around the N-terminal, ubiquitin-like domain of nsp3 (Ubl1). H1 binds tightly to a hydrophobic cleft on the surface of Ubl1, while the polyQ and H2 regions complete a bipartite folding-upon-binding interaction. The interaction results in a massive collapse in the conformational sampling of dimeric N, bringing N2 and N4 in close contact and forming a compact complex with Ubl1. Nsp3 forms the cytosolic extremity of the exit channel of DMVs, suggesting that this essential interaction between viral proteins plays a role in positioning N prior to encapsidation of the nascent RNA, a model supported by the observed accumulation of N in the vicinity of the surface of DMVs (*4*).

The functional importance of N3 has also been highlighted by the accumulation of important mutations, associated with dominant variants of concern (for example: alpha, gamma, omicron ^203^RG-KR, beta ^205^T-I, delta ^203^R-M, omicron ^215^G-C and BA.2.86 ^229^Q-K), in this region (*48*, *49*). Most of these mutations are associated with the SR segment of N3. Indeed mini-replicon studies have measured 10-fold increased mRNA delivery and protein expression for commonly found mutants, and a reverse genetics model revealed 50-fold more virus production for S202R and R203M mutations (*50*). Helix H1 has also been linked to assembly, with a recent study using analytical ultra-centrifugation, circular dichroism combined with molecular dynamics simulation, suggesting that the motif promotes higher order assembly (*51*).

Perhaps most significantly, the SR region of N3 is found to be hyperphosphorylated in infected cells, although is unphosphorylated during viral assembly and is found in the unphosphorylated form when bound to genomic RNA in infectious virions (*52–54*). It has been suggested that hyperphosphorylation or dephosphorylation may play a role as a functional switch in the replication cycle of the virus, either between transcription and replication, or to enable genome packaging or unpackaging (*14*, *55–58*). Phosphorylation has also been shown to affect the compactness of viral ribonucleoprotein (vRNP) complexes (*59*), and concurrent studies to this have shown that binding of the isolated N3 to long RNAs is modulated by phosphorylation (*60*), while fluorescence polarization measured nearly an order of magnitude weaker binding to a 10- nucleotide RNA following hyperphosphorylation (*46*).

N is also known to undergo liquid-liquid phase separation (LLPS) upon mixing with RNA (*26*, *27*, *32–37*), possibly providing compartmentalization of some stage of the viral replication cycle, as has recently been shown for negative-sense single-stranded RNA viruses, such as rabies (*61*) and measles (*62*). Hyperphosphorylation of the SR region (pSR) has also been shown to modulate the nature of membraneless organelles formed from N and RNA (*27*, *34*, *36*, *37*). Although the specific function of SARS CoV 2 viral compartments remains the subject of debate (*63*), it is clear that the storage of high concentrations of N in the vicinity of DMVs (*4*, *6*, *16*, *53*) may provide an efficient mechanism for rapid encapsidation of newly synthesized RNA.

Despite the extensive work described above, little is known about the impact of phosphorylation on the structural and dynamic behaviour of N and its impact on molecular function. The kinase cascade responsible for hyperphosphorylation of N was identified (*58*), implicating sequentially, serine arginine protein kinase 1 (SRPK1), glycogen synthase kinase 3 (GSK-3) and casein kinase 1 (CK1). Here, we use time-dependent NMR spectroscopy, to characterise the conformational and functional implications of hyperphosphorylation of N at atomic resolution. This reveals that transient contacts between H1 and N2, and N4 and N2are abolished upon hyperphosphorylation and replaced by direct binding of pSR to the RNA-binding surface of N2. pSR appears to interact with N2 via the same interface as single stranded RNA, suggesting an auto-inhibitory mechanism involving direct competition with RNA. Notably we demonstrate that phosphorylation by SRPK1 and GSK-3 is sufficient for this inhibition, although not all phosphorylation sites are required and that *in vitro* phosphorylation by non-cognate kinases such as PKA, a commonly used proxy for hyperphosphorylation, does not result in inhibition of RNA binding. In particular time-resolved NMR reveals the precise threshold of the specific phosphorylation pattern necessary to achieve the conformational switching mechanism required to inhibit RNA binding, providing important new insight into the role of post-translational modification in the SARS-CoV-2 replication cycle.

## Results

### Hyperphosphorylation of the disordered SR region of N234

A construct comprising domains N2, N3 and N4 (N234), including the RNA-binding, dimerization and hyperphosphorylation domains, was initially investigated (residues 45-375, lacking only the disordered N- and C-terminal domains). Incubation with SRPK1 resulted in the appearance of two phosphorylation sites, that were assigned to sites pS188 and pS206. Addition of GSK-3 and then CK1 results in phosphorylation of 8 and 4 additional sites respectively, in agreement with recent reports (figure 2). Under physiological conditions, the 31 amino acid SR region thus evolves from a total charge of +6, to +2, -14 and finally -22 over the three phosphorylation steps (resulting in pN234(I), pN234(II) and pN234(III) respectively). Each phosphorylation site was assigned using triple resonance 3D NMR, and real-time NMR was used to compare the rate of phosphorylation of different sites as a function of each kinase. In particular this allows us to identify S176 and S186 that are phosphorylated significantly slower by GSK-3 than S180, S184, S190, S194, S198 and S202. Note that while the ^15^N-^1^H correlation peak of pS176 is resolved, and can therefore be accurately analysed in terms of intensity build-up, pS186 is partially overlapped with pS184 and pS202 and significantly slower than both, complicating further analysis (the non-phosphorylated peak is also strongly overlapped).

**Figure 2.**
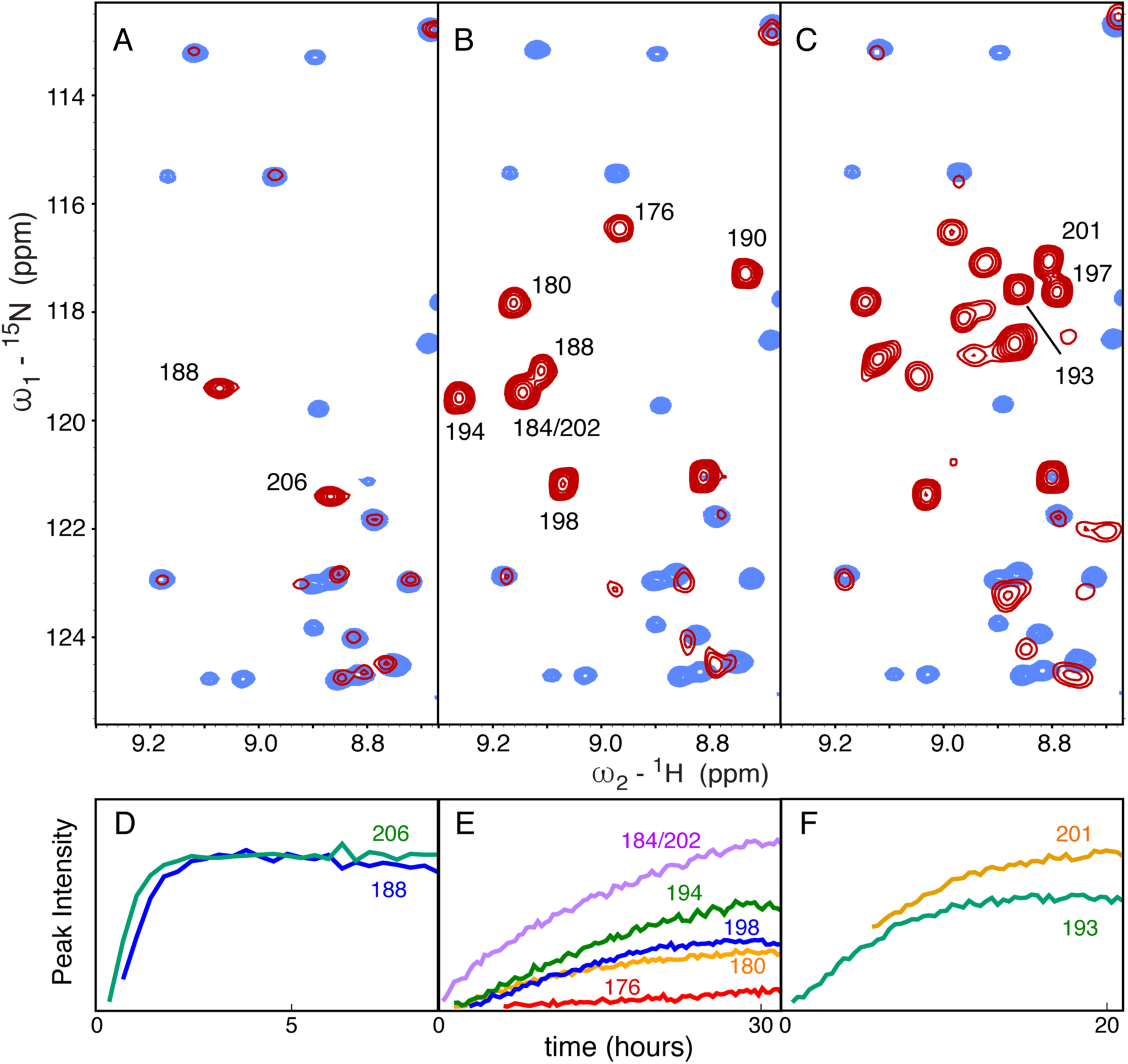
Phosphorylation of N by SRPK1, GSK-3 and CK1. A – Comparison of ^15^N-^1^H HSQC of unphosphorylated (blue) and phosphorylated (red) N234 following incubation with SRPK1 and phosphorylation buffer (referred to as pN234(I)). Two resonances appear that have been assigned to pS188 and pS206. B - Comparison of ^15^N-^1^H HSQC of unphosphorylated (blue) and phosphorylated (red) N234 following incubation of pN234(I) with GSK-3 and phosphorylation buffer (referred to as pN234(II)). Eight additional resonances appear (seven are shown here) that have been assigned to pS176, pS180, pS184, pS186, pS190, pS194, pT198 and pS202. C - Comparison of ^15^N-^1^H HSQC of unphosphorylated (blue) and phosphorylated (red) N234 following incubation of pN234(II) with CK1 and phosphorylation buffer of (referred to as pN234(III)). Four additional resonances appear (three are shown) that have been assigned to pS193, pS197 and pS201. All spectra were buffer exchanged into standard NMR buffer prior to recording these spectra. D-F – Evolution of peak intensity as a function of time for selected resonances that remain sufficiently resolved throughout the kinetic series to accurately measure peak intensities in each step (in phosphorylation buffer). D – pN234(I), E – pN234(II), F – pN234(III).

In order to investigate possible changes in backbone conformation, ^13^C^α^ chemical shifts of pSR were compared to recently proposed random coil chemical shift values (*64*, *65*) for pN234(III), indicating that no significant induction of secondary structure results from hyperphosphorylation (supporting figure 1), as previously proposed for PKA-phosphorylated N (residues 1-209) (*45*).

### Hyperphosphorylation modifies the dynamic behaviour of the disordered SR region

In order to investigate the impact of phosphorylation on the dynamic behaviour of N234 we have measured ^15^N spin relaxation as a function of phosphorylation state (figure 3). This reveals that although N2 appears to maintain its hydrodynamic behaviour, with comparable R_1ρ_ rates in the presence and absence of phosphorylation, the overall correlation time of the SR region increases over the three steps. The increase in pSR is more or less systematic for both pN234(II) and pN234(III), with local peaks observed around residues pS180, pS188 and 202-204. Comparison of transverse relaxation measured from pN123(II) (phosphorylated with SRPK1 and GSK-3) confirms the above observations, and demonstrates that relaxation of the helical region H1 is not significantly impacted by phosphorylation (this helix is too broadened in N234 to measure relaxation with sufficient precision).

**Figure 3.**
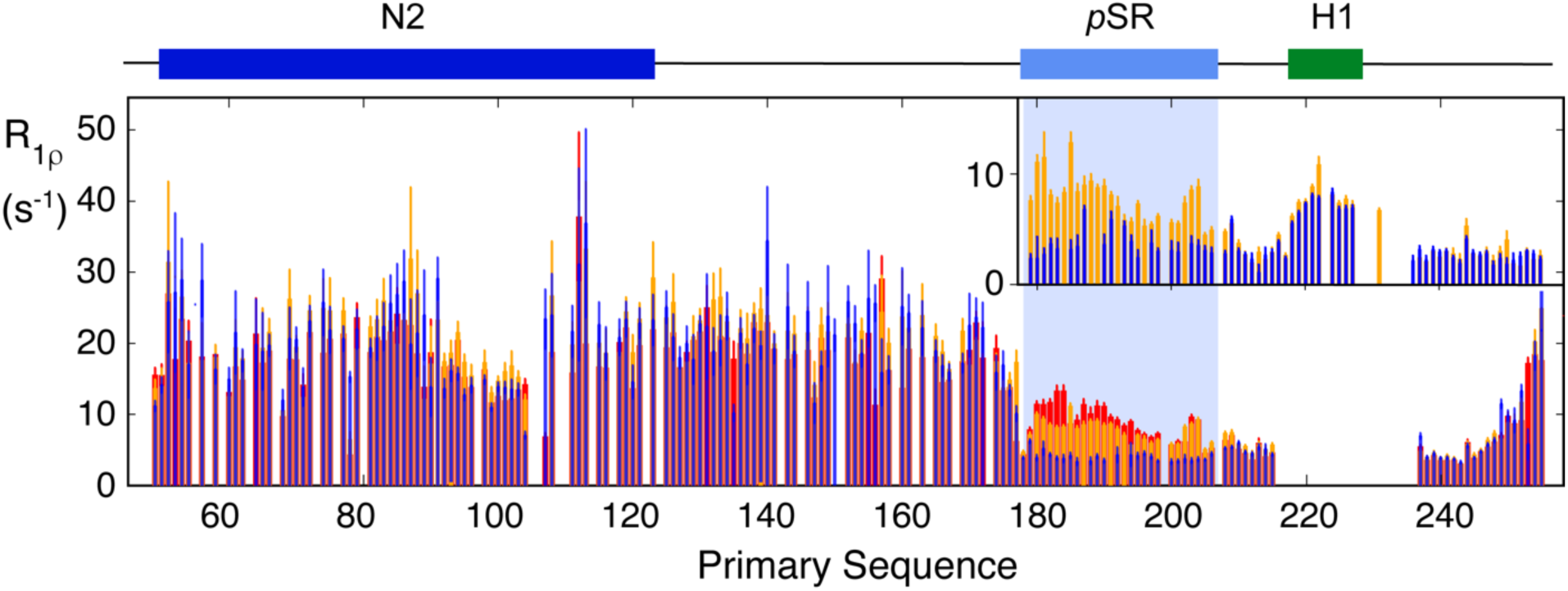
Phosphorylation modifies the dynamic behaviour of N. Main panel - Rotating frame relaxation (R_1ρ_) of N234 at 850 MHz as a function of the phosphorylation state (blue, non-phosphorylated, orange pN234(II), red, pN234(III)). Data from N4 are not shown due to low signal to noise. Inset - Rotating frame relaxation (R_1ρ_) of N123 at 950 MHz (blue, non-phosphorylated N123 and orange pN123(II)). Helix H1 is visible in N123, probably due to the lower molecular weight of this construct, allowing comparison of the dynamic behaviour of this region of N3. This comparison reveals that hyperphosphorylation strongly impacts the dynamic properties of the SR region.

### Interaction of N with RNA from the viral 5’ UTR

We have investigated the RNA binding profile of N234 using a 14 Nt segment (UCUAAACGAACUUU) from the 5’ UTR of SARS-CoV-2 and a 30 Nt polyA. ^15^N-^1^H (CSPs) in N2 measured in N234 (figure 4) resemble those recently published by numerous authors (*25*, *28–31*, *44*), involving the RNA binding finger (residues 90-106), and the surface of the base-plate formed by residues 48-66, 148-160 and 166-174. These observations are confirmed by ^13^C-^1^H methyl chemical shift perturbations (figure 4). Additional shifts are seen throughout the SR region as well as in the boundary between N3 and N4 (residues 244-257), a Lysine-rich region that folds as an α-helix (H2) upon binding nsp3, and forms a detached α-helix in the crystal structure of N4 (*20*, *21*). This region was also shown to interact with RNA in the context of isolated N4 (*43*), as well as in SARS CoV (*39*). Apart from H2, CSPs in N4 are small, except for the solvent exposed C-terminal helix comprising residues 345-357. Notably titration with the two different RNAs resulted in shifts of the same amino acids in N2, and appear upon the same linear CSP trajectories (supporting figure 2), the exact position is dependent on the titration admixture and exchange regime, that differs between RNAs ranging from fast to slow exchange on the NMR timescale, likely related to different affinities (the longer RNA was found to be in slower exchange,).

**Figure 4.**
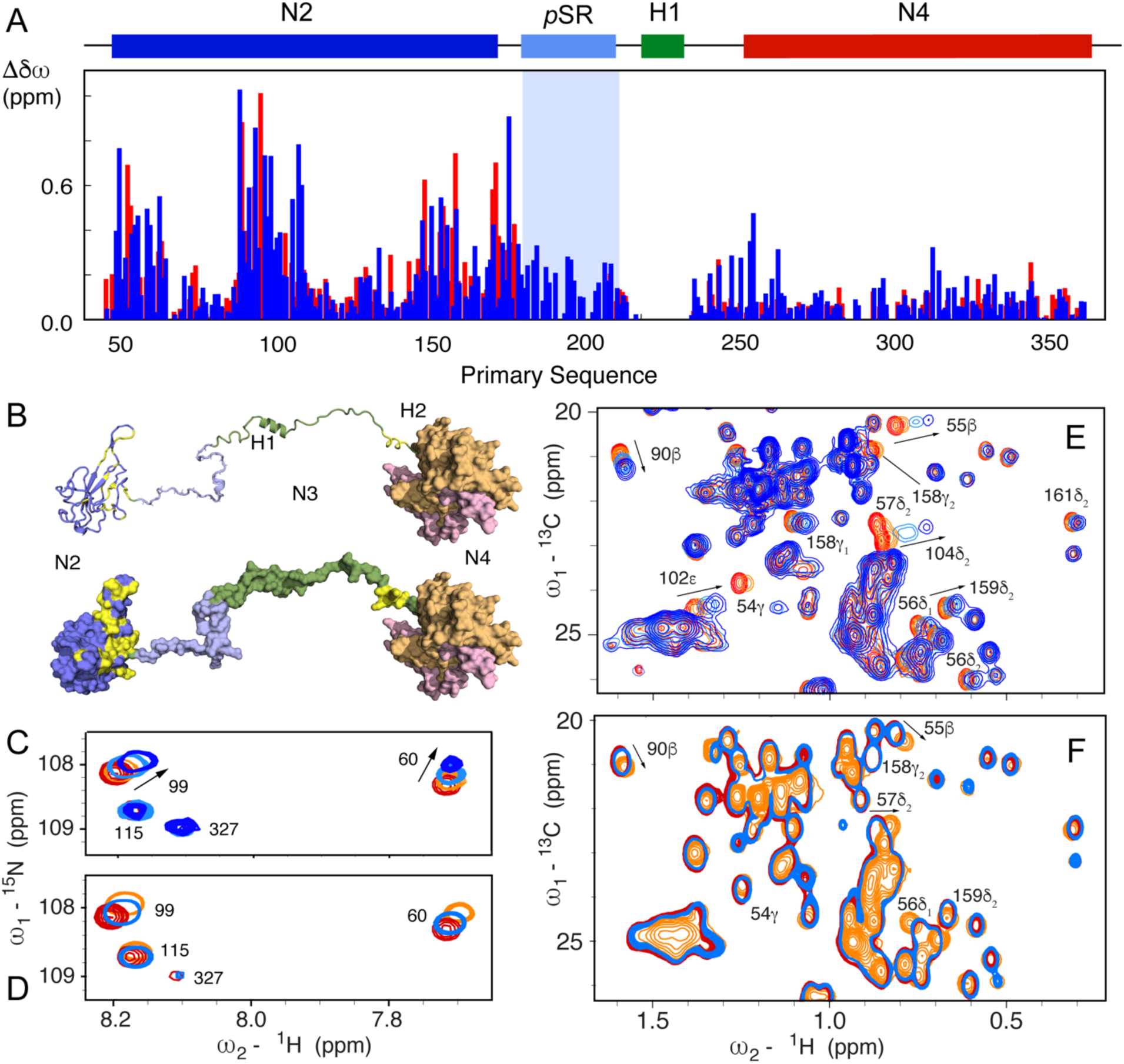
Hyperphosphorylation mimics RNA binding to N2. A – Chemical shift perturbations (CSP) measured in the ^15^N-^1^H TROSY spectrum of N234 following addition of single stranded 14mer RNA (sequence) (dark blue) in comparison with chemical shift differences between non-phosphorylated and SRPK1/GSK-3/CK1 phosphorylated N234 (red) (the extreme shifts from the pSR region due to direct phosphorylation of the residue of interest, are not shown for reasons of dynamic range). CSPs due to RNA binding were scaled by a factor 1.5 for optimal comparison with phosphorylation CSPs. B – Mapping of largest chemical shift differences onto N2 and N3 (yellow), shown in ribbon and surface representation. C – CSPs in ^15^N-^1^H TROSY N234 spectra for selected residues as a function of 14mer single stranded RNA concentration (red – 0%, orange – 25%, light-blue 100%, dark-blue 200%). D - CSPs in ^15^N-^1^H TROSY N234 spectra for the same residues as (C) as a function of phosphorylation state (red – unphosphorylated N234, light blue – pN234(I), orange - pN234(II)). E - CSPs in ^13^C-^1^H HMQC N234 spectra for selected residues as a function of 14mer single stranded RNA concentration (red – 0%, orange – 25%, light-blue 100%, dark-blue 200%). F - CSPs in ^13^C-^1^H HMQC N234 spectra for the same residues as (E) as a function of phosphorylation state (red – unphosphorylated N234, light blue – pN234(I), orange - pN234(II)).

A construct comprising domains N1, N2 and N3 (N123, residues 1-267), including the N-terminal IDR, RNA-binding and hyperphosphorylation domains, was also investigated. The same signature of RNA binding is observed in N2 as in N234 (data not shown), demonstrating that the impact of the dimerization domain on binding of these RNA segments is negligible, or at least too weak to be observed by ^13^C-^1^H and ^15^N-^1^H CSPs. The impact of hyperphosphorylation on the N123 spectrum is also very similar to that observed in N234.

### Hyperphosphorylated SR inhibits N:RNA interaction via the RNA-binding surface of N2

A comparison of the ^15^N-^1^H and ^13^C-^1^H CSPs due to binding of diverse RNAs, and those resulting from sequential phosphorylation by SRPK1, GSK-3 and CK1 (figure 4) reveals remarkable correspondences. While the SRPK1 priming step induces small shifts (see supporting information figure 3), the addition of twelve new phosphorylation sites in pN234(III) results in a very similar distribution of CSPs in N2 compared to those observed as a result of RNA binding (chemical shifts in pN234(II) and pN234(III) show a similar profile, supporting information figure 3). This indicates that hyperphosphorylation induces a change in conformation of N2, probably due to binding of N2 to the pSR region, that mirrors the impact of RNA binding. We note that although the CSPs are distributed similarly, they can be different in sign to those observed upon binding RNA, demonstrating that while the same sites on the surface of N2 are affected, the bound state chemical shifts (RNA-bound and pN3-bound) are different (figure 4).

We then tested the RNA-binding properties of the different forms of pN234. While phosphorylation due to SRPK1 has no observable effect on RNA binding, the additional phosphorylation sites resulting from incubation with GSK-3 and CK1 result in complete abrogation of RNA binding (figure 5A). This was found to be the case for both RNA molecules that were tested. Both pN234(II) and pN234(III) fail to bind RNA, demonstrating that phosphorylation by SRPK1/GSK-3 is necessary and sufficient. We note that it has been previously observed that phosphorylation by SRPK1 is an essential step for phosphorylation by GSK-3 (*58*).

**Figure 5.**
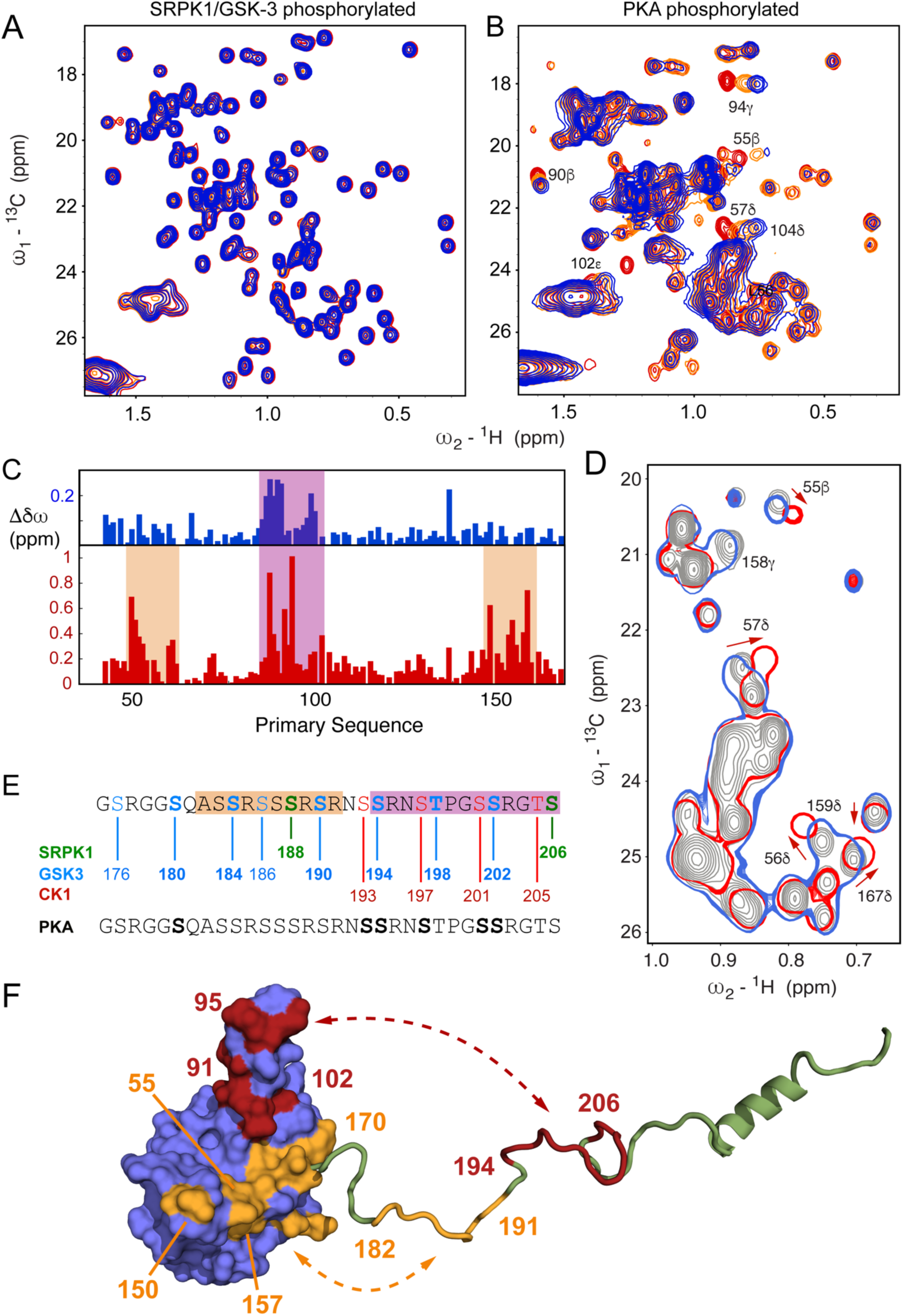
Hyperphosphorylation by SRPK1, GSK-3 and CK1 inhibits RNA binding. A – Chemical shifts in ^13^C-^1^H HMQC spectra of SRPK1/GSK-3 phosphorylated N234 (pN234(II)), for selected residues as a function of 14mer single stranded RNA concentration (red – 0%, orange – 25%, light-blue 100%, dark-blue 200%). No significant shifts were observed (compare to figure 4E). B - Chemical shifts in ^13^C-^1^H HMQC spectra of N234 phosphorylated by PKA, for selected residues as a function of 14mer single stranded RNA concentration (red – 0%, orange – 25%, light-blue 100%, dark-blue 200%). Shifts are highly comparable with CSPs measured in unphosphorylated N234 (compare to figure 4E). C – CSPs (blue) in ^15^N-^1^H TROSY of N234 phosphorylated by PKA, and (red) pN234(II). D – Section of the ^13^C-^1^H HMQC spectrum of unphosphorylated N234 (grey), with N234 phosphorylated by PKA (blue) and pN234(II) (red). E – Sequence of the SR region showing *in vitro* phosphorylation sites (bold) following incubation with PKA (grey/black) and, sequentially SRPK1 (green), GSK-3 (blue) and CK1 (red). F – Sketch illustrating putative differential binding sites of the N-terminal (182-191) and C-terminal (194-206) regions of pSR on the surface of N2. Red shading on N2 indicates chemical shifts that are induced by both PKA and SRPK1/GSK-3 phosphorylation, while orange indicates regions that are only shifted significantly by SRPK1/GSK-3 phosphorylation.

The broadly phosphorylating protein kinase A (PKA) has been exploited to phosphorylate N in a single step, including via co-expression with N (*45*), and notably to investigate the impact on binding of the host factor 14-3-3 (*66*). Incubation of N with PKA *in vitro* results in phosphorylation of six sites in the SR region of N234 (supporting figure 4) whose backbone resonances were assigned, namely pS180, pS193, pS194, pS197, pS201 and pS202, although broader phosphorylation was observed by co-expression with PKA (*45*, *66*). This 6-fold phosphorylated form of N234 nevertheless maintains its binding to RNA (figure 5B). The specific pattern resulting from SRPK1/GSK-3 phosphorylation is therefore critical for abrogating RNA-binding to N234, while N234 phosphorylated by PKA *in vitro* is unable to reproduce this functional phenotype.

The possible origin of this differentiation is revealed by comparison of ^15^N-^1^H and ^13^C-^1^H chemical shift perturbations due to PKA phosphorylation of N234 compared to SRPK1/GSK-3 phosphorylation (figure 5 C,D). Although much smaller shifts are seen in general, the main interacting regions appear to involve the RNA-binding finger (90–106). CSPs involving the base-plate (48–66, 148–174) are considerably smaller. Notably, the region of pSR situated between positions 182 and 191 comprises four phosphorylation sites in pN234(II), but is unmodified by *in vitro* PKA-phosphorylation (figure 5E). Taken together, these observations are consistent with a model whereby phosphorylation of the N-terminal strand of pSR (residues 182-191) is responsible for interaction with the base-plate, while phosphorylation of the C-terminal strand (residues 194-206), present in both pN234(II) and PKA-phosphorylated N234, would be responsible for binding the RNA-binding finger (figure 5F). It is tempting to speculate that occupation of both interaction sites is required to abrogate interaction with RNA. Note that the specific differences in phosphorylation pattern of residues 194-206 may also contribute to stronger and more efficient binding of N2.

### Hyperphosphorylated SR sites specifically abrogate RNA binding

In order to further identify the specific phosphorylation steps that are essential for abrogating RNA binding to pN234(II), modulation of the RNA binding equilibrium was investigated during the phosphorylation process. Having established that the minimum phosphorylation level for abolishing RNA interaction occurs as a result of phosphorylation by SRPK1 and GSK-3, the 14-mer RNA was added to pN234(I), to which GSK-3 and ATP were added. Signature chemical shifts reporting on single stranded 14mer RNA binding were followed using ^13^C-^1^H HMQC methyl spectra, and the level of phosphorylation followed by measuring intensities of interleaved ^15^N-^1^H SOFAST HSQC, as described above. This allows us to follow the population shift between free and bound RNA by measuring sites that shift upon RNA binding (for example methyl groups of I94 and M101), and compare this with the rate of appearance of the 8 sites that are phosphorylated by GSK-3, where the covalent nature of the modification provides a direct measure of the extent of the phosphorylation.

Increases in intensity of resolved phosphorylation peaks are compared to the average chemical shift evolution of the methyl groups (figure 6A,B). This comparison reveals that while S180 phosphorylates faster than the observed shift reporting on the RNA binding equilibrium, a number of sites show rates of phosphorylation that are very similar to the rate of RNA unbinding (S184, S194, T198, S202), and two sites have significantly slower phosphorylation rates (S176 and S186), suggesting that the latter two sites are not essential for the inhibition of RNA binding, at least under the conditions we have tested *in vitro*. The experiment was repeated using 30-mer polyA RNA. The evolution of the chemical shift of Met 101ε is compared to the intensity of the phosphorylation peak of pS194, demonstrating that inhibition of RNA binding is again effective in the case of this longer RNA molecule (figure 6 C,D).

**Figure 6.**
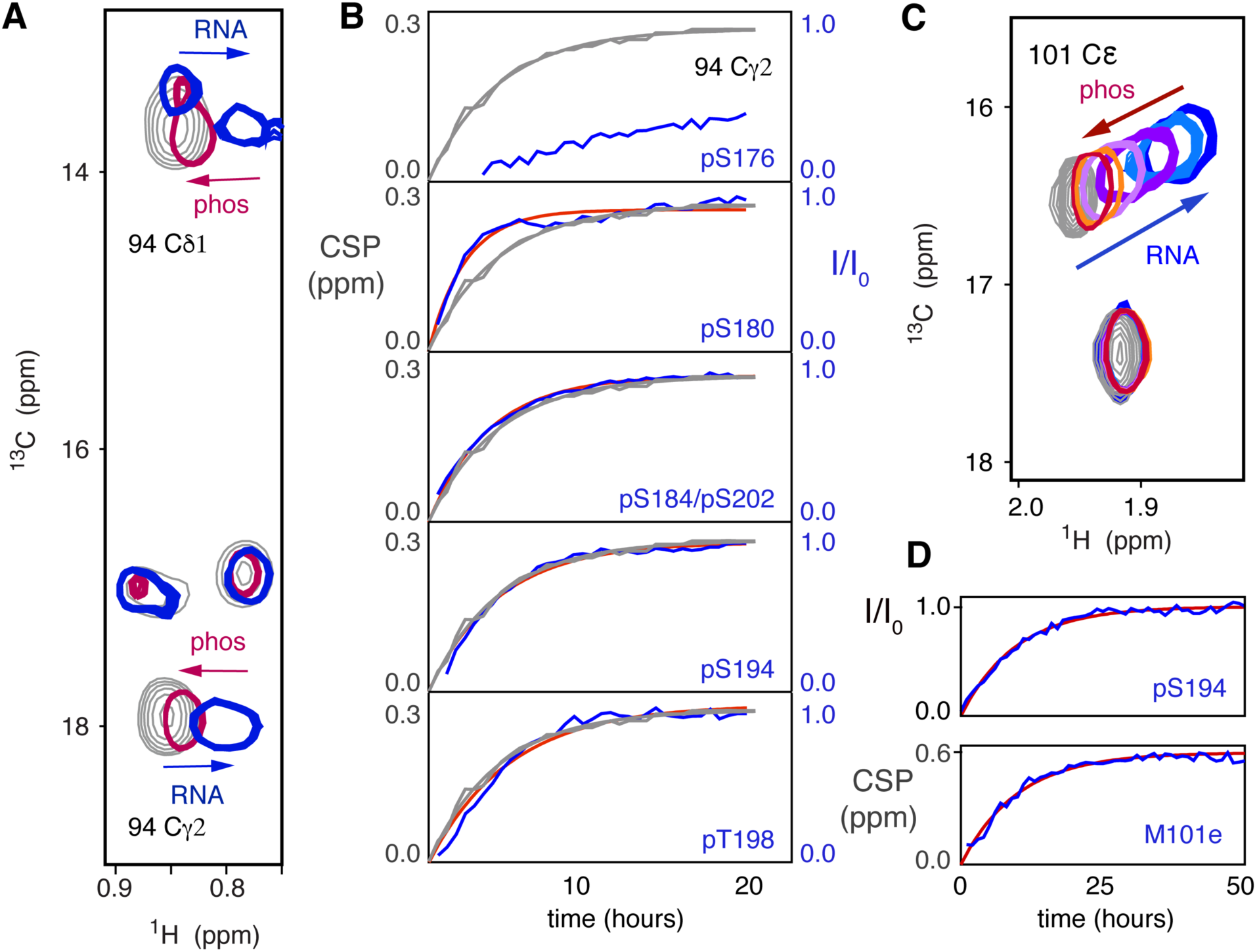
Real-time observation of RNA binding inhibition by SR phosphorylation. A – CSPs in methyl ^13^C-^1^H HMQC of pN234(I) (specifically 94Cψ2, showing impact of 14mer RNA binding (grey, 0μM, blue 200μM), and subsequent shift of the same peaks back towards the unbound form during incubation with GSK-3 (red). B – Comparison of time course of intensity increase in the ^15^N-^1^H HSQC phosphorylation peaks from *p*SR (blue, experimental, red, single exponential fit) and the chemical shift perturbation of 94Cψ2 (grey, experimental and fitted). Associated time constants of (0.50±0.07)h (pS180), (0.30±0.04)h (pS184/pS202), (0.26±0.07)h (pS194) and (0.24±0.05)h (pT198) compared with (0.24±0.03)h and (0.26±0.03)h for 94Cψ2 and 94C81. pS176 and pS186 (not shown) were too slow to be accurately fitted. C – CSPs in methyl ^13^C-^1^H HMQC of pN234(I) (M101Cχ) showing impact of 30mer polyA RNA binding (grey, 0μM, blue 120μM), and subsequent shift back towards unbound form during incubation with GSK-3 (light blue to red). D – Comparison of (top) the time course of increase in intensity in the ^15^N-^1^H HSQC pS194 peak (blue - experimental, red - fitted to a single exponential) and (bottom) the shift of methyl ^13^C-^1^H CSP of 101Cχ back from RNA-bound to unbound form during incubation with GSK-3 as shown in panel C (blue - experimental, red - single exponential fit). Associated time-constants : (0.10±0.04)h pS194 and (0.099±0.030)h for 101Cχ.

### Modulation of long-range order in N due to hyperphosphorylation

In order to develop further insight into the origin of CSPs observed upon phosphorylation, and to further investigate long-range interactions within N234, we have measured paramagnetic relaxation enhancements (PREs) of cysteines mutants of N234, labelled with 4-maleimido-TEMPO. Positions for cysteine mutation were selected on N2 (175) and N3 (210) (figure 7).

**Figure 7.**
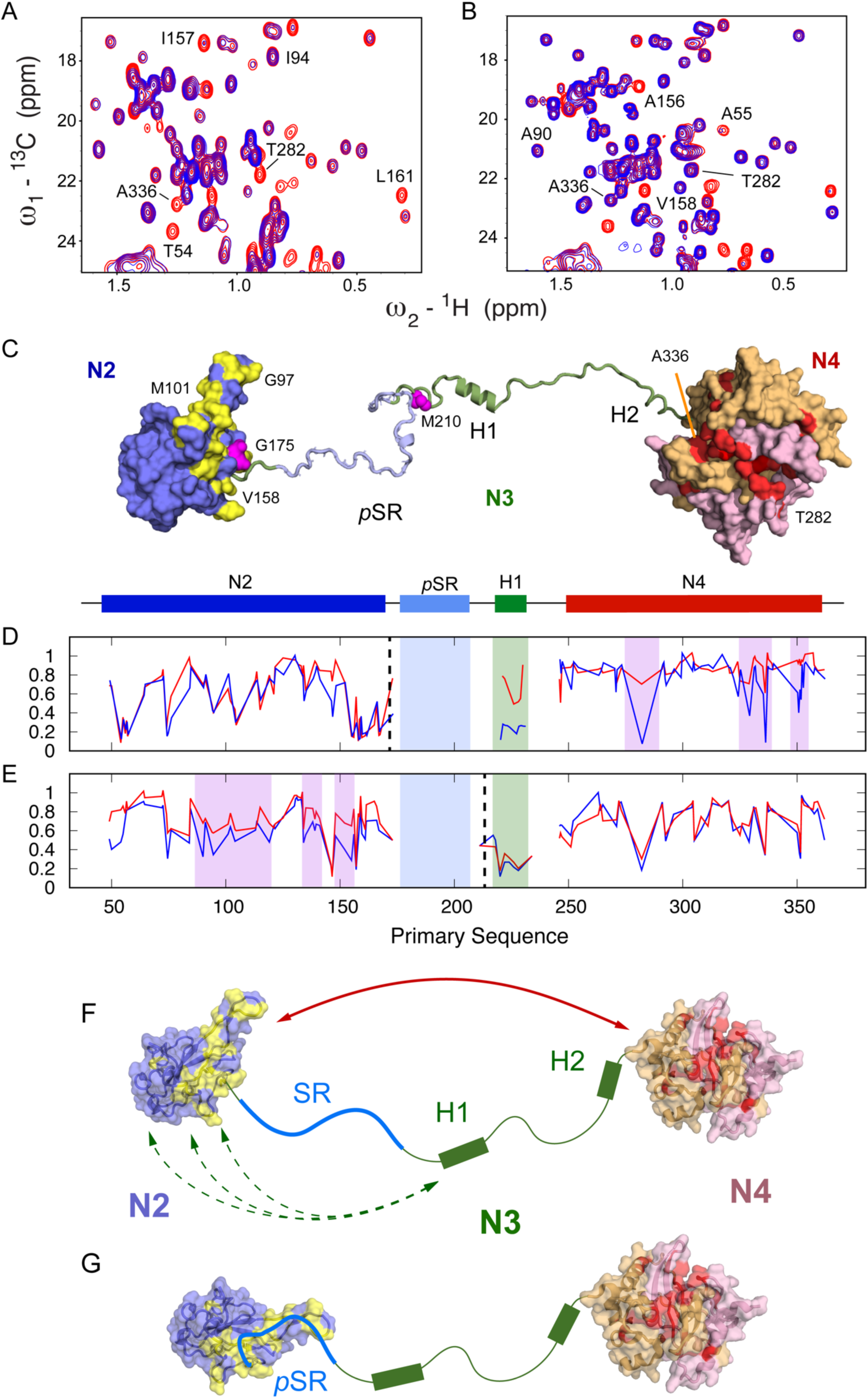
Modulation of paramagnetic relaxation enhancement due to hyperphosphorylation. A – Comparison of ^13^C-^1^H HMQC in oxidised (blue) and reduced (red) forms of TEMPO-labelled N234 (G175C). B – Comparison of ^13^C-^1^H HMQC in oxidised (blue) and reduced (red) forms of TEMPO-labelled pN234(II) (G175C). Strong interdomain PREs highlighted in panel A (282 and 336) are considerably weakened due to phosphorylation. C – Surface representation indicating selected effects on long-range order in N234. Red – residues on hydrophobic surface of N4 that broaden in the presence of TEMPO label (175), that is weakened upon phosphorylation. Beige (N2) – residues on RNA binding finger that broaden in presence of TEMPO label (210), also weakened upon phosphorylation. Yellow – residues on N2 that shift upon hyperphosphorylation. Sites of the TEMPO labels are shown in magenta. D-E. Intensity ratios between paramagnetic and diamagnetic forms of the protein. Blue – free N234, red – pN234(II). Dashed lines indicate position of the TEMPO label (175 and 210). F-G. In the free, non-phosphorylated form (A) long-range interactions between N2 and N4 and between H1 and N2 are revealed from PREs. Upon hyperphosphorylation interactions are weakened (B) or completely suppressed. CSPs on N2 (yellow) (see panel C).

In non-phosphorylated N234, a hydrophobic region of N4, covering residues 333-340, 350 and 282, is broadened by the label positioned at residue 175 (figure 7), indicating that this region is involved in a long-range contact between the two folded domains. In addition, the linker region, in particular the peaks corresponding to the hydrophobic groups of H1, is strongly broadened. The suggestion that H1 interacts transiently with N2 is supported by data from the paramagnetic label at position 210, which induces broadening in N2, both in the RNA-binding finger (residues 85-105) and the C-terminal region (residues 145-170) and is supported by recent similar observations using PREs from neighbouring sites in a shorter construct (*30*). Broadening is also observed on the hydrophobic patch on N4 due to the label at position 210.

^15^N chemical shifts of N2 were compared in the presence and absence of N1, N3 and N4. Small differences are observed between free N2 and the longer constructs (supporting figure 5), clustering mainly around the base-plate residues 60-70 and 160-170, located on a single continuous face of N2, suggesting that the weak interactions with N4 observed from PREs involve this surface.

In order to probe the impact of phosphorylation on long range order, two forms of the TEMPO labelled protein (positions 175 and 210) were also hyperphosphorylated (pN234(III)) and PREs again measured (figure 7). Notably the broadening profile of the probe on N2 reveals significant and systematic abrogation of the long-range contacts between N2 and the hydrophobic patch on N4. This surface participates in pSR interactions with N2, suggesting that the long-range contacts between N2 and N4 are removed due to the presence of bound pSR on the surface of N2. In addition, the enhanced broadening in H1 due to the label on N2 is weakened due to hyperphosphorylation, indicating that interactions of H1 with the surface of N2 are also abolished. This is supported by abrogation of PREs due to the 210 label that implicate the RNA binding finger (residues 80-110). The PRE of N4 due to 210 is not strongly impacted by phosphorylation.

The relevance of this abrogation of long-range contacts may be related to assembly or disassembly of nucleocapsids. In this respect we note that we used negative staining electron microscopy to reveal the presence of cage-like particles upon addition of RNA (supporting figure 6). Such structural elements have been previously identified as substructures of the viral nucleocapsid (*15*, *16*). The presence of these assemblies was not observed when RNA was added to hyperphosphorylated N234, confirming recent observations (*46*). In the absence of atomic resolution structures of assembles nucleocapsids it is not yet clear whether this impact on the formation of larger assemblies is related to the effect of phosphorylation on long-range contacts in N.

## Discussion

The SR region of SARS-CoV-2 nucleoprotein has been shown to be hyperphosphorylated in infected cells, and unphosphorylated during viral assembly and when bound to genomic RNA (*52*, *53*). The level of phosphorylation of N is thought to act as a functional switch in the replication cycle, or in genome packaging or unpackaging (*14*, *55–58*). Phosphorylation has also been shown to affect the compactness of viral ribonucleoprotein (vRNP) complexes (*59*), and concurrent studies have shown that binding of the isolated N3 to long RNAs is modulated by phosphorylation (*60*), while fluorescence polarization measured significantly weaker binding to a 10-nucleotide RNA following hyperphosphorylation (*46*). A recent study proposed the sequential phosphorylation of N by host kinases SRPK1, GSK-3 and CK1 (*58*). Inhibition of SRPK1 was shown to reduce replication, while blocking the activity of GSK-3 reduced viral replication both in cells, and lowering infection in patients (*67*).

It is not yet clear where phosphorylation plays a role in the viral cycle of SARS-CoV-2. While nucleocapsids present in the virion are not phosphorylated, N is found in the hyperphosphorylated form in the cytosol of infected cells. Phosphorylation is thought play a role in unpackaging (following infection) and packaging (prior to viral assembly). Mass photometry and electron microscopy were recently combined to reveal that phosphorylated N forms less compact viral RNPs (*59*).

Despite this intense interest, little is currently known about the impact of hyperphosphorylation on the structural, dynamic and functional behaviour of N. This study exploits the site-specific information provided by high resolution NMR to describe the effects on local and long-range structure as a function of the proposed phosphorylation cycle, as well as the implications of sequential post-translational modifications on molecular function.

Our analysis substantiates the reported impact of phosphorylation by the previously identified putative triplet of host kinases of SRPK1, followed by GSK-3 and CK1 (*57*, *58*). This confirms that the proposed sites are indeed phosphorylated (S206 and S188 by SRPK1, followed by S176, S180, S184, S186, S190, S194, T198 and S202 by GSK-3 and S193, S197, S201 and S205 by CK1), and that the order of phosphorylation by GSK-3 reveals that S180 is phosphorylated before near simultaneous phosphorylation of S184, S190, S194, T198 and S202, followed by slower phosphorylation of S176 and S186. These observations are confirmed both by monitoring the peaks reporting on the amide group of the directly phosphorylated amino acid, as well as of the immediately neighbouring amino acids (supporting figure 7).

Phosphorylation affects the dynamic properties of N locally, in the SR hyperphosphorylation domain. Rotating frame relaxation rates increases (figure 3) as a function of phosphorylation state. The progressive increase with increasing levels of phosphorylation may result from rigidification of local structure in pSR, due to the increased presence of bulky phosphate groups, and/or from interaction with another domain of N. The latter appears to be the case, with hyperphosphorylation inducing a highly similar profile of chemical shift perturbations compared to those observed due to RNA binding to N234 (figure 4). These changes are predominantly evident in N2, but also in the C-terminus of N3, comprising a Lysine-rich helical region (H2) that is formed in the presence of N4, and that folds into a helical conformation when N3 binds to nsp3 (*68*). It therefore appears that pSR binds via the same interface on N2 as that exploited by RNA. Indeed, while phosphorylation by SRPK1 has no notable impact on the N:RNA interaction, phosphorylation by GSK-3 inhibits binding completely. Using time-resolved NMR, at residue-specific and atomic resolution, identifies which phosphorylation sites are essential for this inhibition, including most sites modified by GSK-3, with the exception of S176 and S186. The necessary phosphorylation pattern for inhibition of RNA binding is thereby revealed as a regularly spaced charge distribution with respect to the primary sequence: S180, S184, S188, S190, S194, T198, S202, S206, underlining the probable role of electrostatics in this inhibition. These sites are all known to be highly conserved within the SR region.

Progressive phosphorylation of SR therefore appears to act as a conformational and functional switch for SARS-CoV-2 nucleoprotein, with time-resolved NMR providing crucial insight into the role of each phosphorylation site in the functional mechanism. It is tempting to suggest that recruitment of host kinases may thereby participate in releasing genomic RNA from encapsidation, upon unpackaging from the infecting virion after cell-entry. The observation that SRPK1 and GSK-3 are apparently sufficient to achieve this switch also raises the question of the role of CK1 phosphorylation in viral function.

More insight into this putative interaction between pSR and the RNA binding interface of N2 is provided by comparison with the impact of *in vitro* phosphorylation by PKA. Under our experimental conditions, this results in phosphorylation of S180, S193, S194, S197, S201 and S202, generating an additional 12 negative charges in the SR region, compared to the 16 that are required to abrogate RNA binding upon SRPK1/GSK-3 phosphorylation. Crucially however, phosphorylation by PKA has no measurable impact on RNA binding, suggesting that the pattern resulting from the physiologically relevant kinases is indeed specific for inhibition, either by binding tighter to the surface of N2, by occupying more RNA binding sites on N2, or by ensuring a uniform electrostatic screening of the RNA-binding surface. A recent study provided evidence that phosphorylation of N(1–209) with PKA resulted in chemical shifts of the β-hairpin (91–105) of N2 (*45*). Our measurements confirm this observation. Indeed, differences in the chemical shifts induced by phosphorylation with PKA and SRPK1/GSK-3 suggest that the PKA phosphorylated SR does not interact with the base-plate regions (residues 48-66, 148-174) as is the case for SRPK1/GSK-3 phosphorylated SR. Phosphorylated SR interacts with the RNA-binding finger in both cases, although inducing much smaller CSPs in the case of PKA phosphorylation. This allows us to propose distinct interaction sites between the C- and N-terminal strands of pSR with the RNA-binding finger and baseplate (48–66, 148–174) of N2 respectively (figure 5). This occupation of additional RNA-binding sites on N2 by SRPK1/GSK-3 phosphorylated N234 may explain why the conformational switch necessary to abrogate RNA binding is effective in this case and not in the case of PKA phosphorylation.

Paramagnetic relaxation NMR was used to characterise the modulation of long-range order induced by phosphorylation. In the free, unmodified protein, helix H1 interacts transiently with the surface of N2, possibly mediated by hydrophobic interactions between the Leucine-rich motif at the N-terminus of H1 and hydrophobic sites on N2, and specifically involving the RNA-binding finger (85–105) and the (145–170) region, also implicated in RNA binding. Helix H1 has previously been implicated in binding to N2 (*30*), indeed we find some evidence of abrogation of transient contacts with N2 as a result of hyperphosphorylation, although the negligible effect of phosphorylation on transverse relaxation of H1, and the lack of chemical shift perturbation in N2 in the presence or absence of N3, suggests that such interactions must be weakly populated.

Stronger interactions are also observed between N2 and N4, specifically involving a hydrophobic patch on one side of the dimeric interface of N4, comprising 333-340, 350 and 282, and the base-plate of N2, suggesting a highly dynamic equilibrium between different states, as previously proposed on the basis of small angle scattering (*68*). This equilibrium is modified upon hyperphosphorylation, probably because the N2 interaction surface is occupied by the pSR region. Clear differences can be identified, with notable weakening, even disappearance, of the interactions implicating the hydrophobic patch on N4. This abrogation of contacts within a single N dimer may correlate with the macroscopic observation of looser nucleocapsid structures derived from electron microscopy (*59*), or the reported modulation of the dynamics of membraneless organelles that can be formed upon mixing N and RNA under certain conditions (*26*, *34*). Indeed, we note that under our experimental conditions cage-like structures are evident by negative staining electron microscopy upon mixing RNA with non-phosphorylated N234, but that such structures are absent for hyperphosphorylated N234/RNA mixtures.

Phosphorylation of disordered peptides is known to facilitate rapid switching of biomolecular function of many physiological processes (*69–71*). Progressive phosphorylation of newly unpackaged N by host-kinases likely maintains the protein in a form that is incapable of binding RNA, via an autoinhibitory strategy that protects from non-specific binding to cytosolic host RNA, as well as premature binding to viral genomic RNA. The clear identification of a threshold of phosphorylation, achieved during GSK-3 phosphorylation, identifies specific molecular characteristics of this switching mechanism, a process that establishes phosphorylated N in an RNA-unbound form that may participate in additional functional activity such as regulation of transcription (*14*). Newly expressed N is apparently not phosphorylated prior to encapsidation, assembly and packaging, suggesting that a strategy for inhibition of phosphorylation must be available. It has been suggested that replication occurs within membraneless organelles formed by liquid-liquid phase separation of N (*6*, *26*, *27*, *32–35*, *37*, *63*, *72*, *73*), and formation of organelles may provide some kind of protection from host kinases. Assuming encapsidation of newly synthesized RNA can occur within such organelles, as is the case for numerous negative strand RNA viruses (*61*, *62*, *74–76*), such a mechanism would provide an efficient means to escape from host kinase phosphorylation and facilitate efficient encapsidation of genomic RNA by unphosphorylated nucleoprotein.

## Conclusion

In conclusion, we have used NMR spectroscopy to describe the impact of phosphorylation on the serine-arginine rich region of the central disordered domain of SARS-CoV 2 nucleoprotein, a region that is hyperphosphorylated in infected cells. The proposed phosphorylation scheme, comprising SRPK1, GSK-3 and CK1, is shown to abrogate binding of RNA. The phosphorylated SR domain interacts with the RNA binding domain of N via the same interface as single stranded RNA, suggesting an auto-inhibitory mechanism involving direct competition with RNA. Long-range interactions between the folded domains of a single N protomer are also abrogated by phosphorylation, an observation potentially related to packaging or unpackaging of the viral genome. It seems likely that the observed impact of phosphorylation of SARS-CoV-2 N on RNA binding reports on a switching mechanism that inhibits non-specific binding of N to viral, or host RNA. The ability to determine the precise threshold of the highly specific phosphorylation pattern required to inhibit RNA binding provides crucial new insight into the role of post-translational modification in the viral cycle and identifies potential new opportunities for inhibitory strategies for this class of viruses.

## Materials and Methods

### Protein expression and purification

All proteins (N234 47-364, and N123 1-267) were expressed in *E. coli* and purified as described previously. Single-point mutations of N234 (G175C and M210C) were made in-house by site-directed mutagenesis. 14mer RNA (UCUAAACGAACUUU) and 30mer polyA were purchased from Integrated DNA Technologies (IDT), San Diego.

### Phosphorylation of N123 and N234

N123, N234, and G175C and M210C mutations of N234 were incubated with SRPK1 (Sigma-Aldrich), GSK-3 (Promega), CK1 (Promega), or PKA (Promega) kinases in phosphorylation buffer: 50 mM Na-phosphate (pH 6.5), 250 mM NaCl, 2mM DTT, supplemented with 10 mM ATP, and 10 mM MgCl2 at 25°C. For phosphorylation cascade reaction kinases (SRPK1, GSK-3, CK1) were added to the sample sequentially or simultaneously.

### Nuclear magnetic resonance experiments

Unless stated otherwise, all NMR experiments were carried out in 50 mM Na-phosphate (pH 6.5), 250 mM NaCl, 2mM DTT or 50 mM Na-phosphate (pH 6.5), 250 mM NaCl, 1mM TCEP at 25°C. Experiments were acquired on Bruker spectrometers with ^1^H frequencies of 600, 700, 850 and 950 MHz. Spectra were processed with NMRPipe (*77*) and visualized using NMRFAM-Sparky (*78*).

^15^N R_1ρ_ relaxation was measured at 298K and a ^1^H frequency of 850 MHz (N234 and pN234) and 950 MHz (N123 and pN123) using a spin lock of 1.5 kHz (*79*). Relaxation delays of 1, 10, 30, 30, 70, 120, 200 ms were used for ^15^N R_1ρ_ of N234, and 1, 10, 20, 50, 50, 90, 140, 180, 250 ms were used for ^15^N R_1ρ_ of N123 and included repetition of one delay. Relaxation rates were fitted using in-house software and errors estimated with noise-based Monte Carlo simulation.

#### Resonance assignment

Backbone assignment of domains N1 to N4 in the non-phosphorylated forms of N123 and N234 have been recently published (*17*, *80*). Assignment of the phosphorylated forms was performed using BEST-type triple resonance experiments (*81*). Sidechain assignment of N2 (*25*) and N4 from SARS CoV (*39*) have been published and served as references to complete the assignment of the methyl region of ^13^C-^1^H HMQC of N234. ^13^C^α^ chemical shifts were compared to random coil values using the program SSP (*82*). Chemical shift perturbations (CSPs) were calculated using the following expression (*83*)

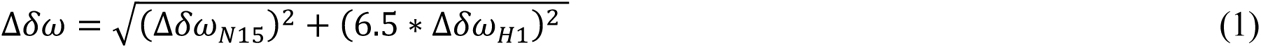

#### PRE experiments

PRE effects were measured from the peak intensity ratios by comparing ^15^N-HSQC and HMQC 2D spectra recorded on samples labelled with TEMPO and samples reduced by the addition of 2 mM ascorbic acid. Briefly, purified cysteine mutants were reduced with 10 mM of DTT at 4 °C for 12 hours and dialysed into 50 mM phosphate buffer pH 7.0 containing 250 mM NaCl without DTT. 10-fold molar excess of 4-Maleimido-TEMPO (Sigma-Aldrich) dissolved in DMSO was added to a 100μM stock of the reduced cysteine mutants. The reaction was incubated for 2 hours at room temperature and then left overnight at 4 °C. In order to eliminate the excess of TEMPO, the samples were dialyzed into NMR buffer.

#### Competition experiments

SRPK1-phosphorylated N234 in phosphorylation buffer were supplemented with 200% of 14mer or 120% of polyA30 RNA. Kinases were added directly to the NMR tube, followed by interleaved, sequential ^15^N and ^13^C correlation spectra in the presence of GSK-3 kinase recorded using BEST-TROSY (*84*) and SOFAST-HMQC (*85*).

#### Electron microscopy

The particles used for negative-stain electron microscopy were obtained by adding 10 μM 14mer RNA to 10 μM N234 non-phosphorylated, or phosphorylated with the three kinases, in a final volume of 20μl. Samples were incubated for 1 hour at room temperature, then applied to the clean side of carbon on mica (carbon/mica interface), and stained with 2% sodium silicotungstate (pH 7.0). Micrographs were taken with a T12 FEI microscope at 120 kV and a magnification of 30,000.

## Funding

This work was supported by the European Research Council Advanced Grant DynamicAssemblies under the European Union’s Horizon 2020 research and innovation program (grant agreement number 835161) (M. B.).

This work used the platforms of the Grenoble Instruct-ERIC center (ISBG ; UAR 3518 CNRS-CEA-UGA-EMBL) within the Grenoble Partnership for Structural Biology (PSB), supported by FRISBI (ANR-10-INBS-0005-02) and GRAL, financed within the University Grenoble Alpes graduate school (Ecoles Universitaires de Recherche) CBH-EUR-GS (ANR-17-EURE-0003).

ACZ received funding from the European Union’s Horizon 2020 research and innovation programme under the Marie Skłodowska-Curie grant agreement No. 796490 and HFSP postdoctoral HFSP fellowship LT001544/2017.

Financial support from the IR-RMN-THC Fr3050 CNRS for conducting the research is gratefully acknowledged.

IBS acknowledges integration into the Interdisciplinary Research Institute of Grenoble (IRIG, CEA).

## Author contributions

MBa, ACZ and MB conceived of the project and planned its execution. MBa, ACZ, LMB, SG, EM, NS and JT ran NMR experiments. DM, MBa, ACZ, SG, LMB prepared protein samples. MBa, ACZ, LMB, SG, JT, TH and MB analysed spectra. EM ran and analysed electron microscopy experiments. MB supervised the study and wrote the article. All authors contributed to the final version.

## Competing interests

All authors declare they have no competing interests.

## Data and Materials Availability

All data needed to evaluate the conclusions in the paper are present in the paper and/or the Supplementary Materials.

## Supplementary Materials

**Supporting Figure 1.**
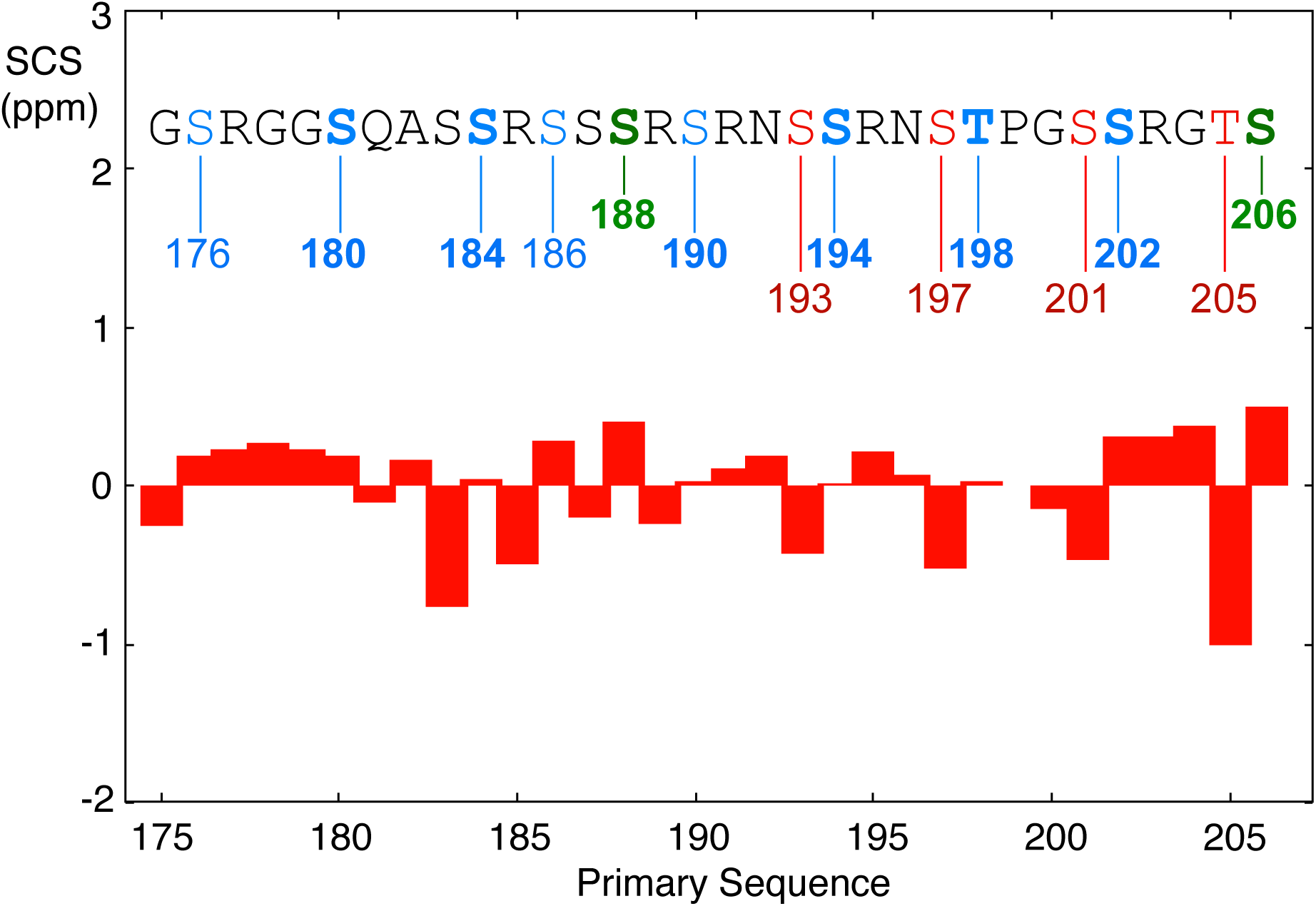
Secondary structural analysis of the pSR region. Ca shifts were compared to recently proposed random coil shifts for phosphorylated peptides (*64*). The position of phosphorylation sites due to the different kinases are shown above in green (SRPK1), blue (GSK-3) and red (CK1).

**Supporting Figure 2.**
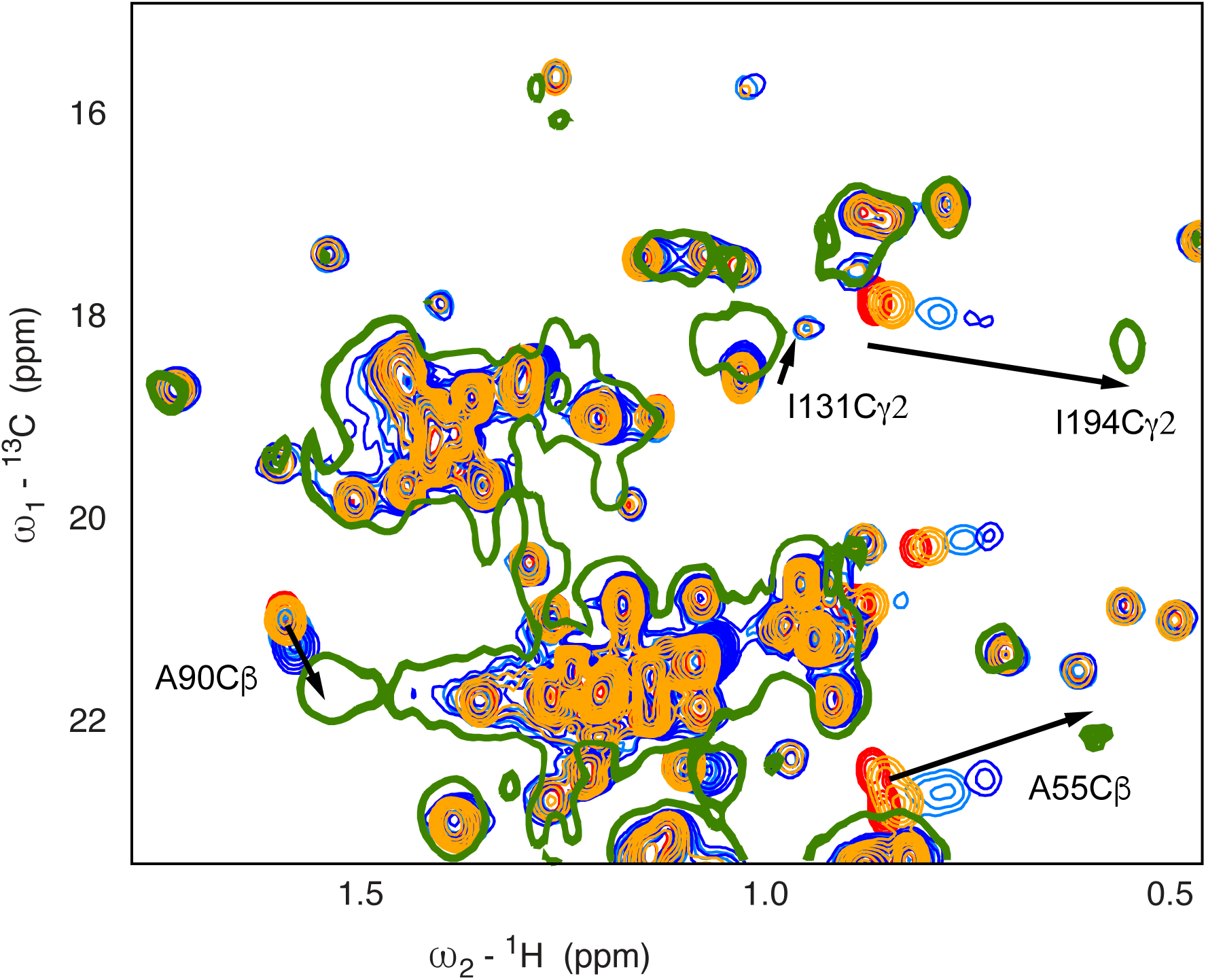
Comparison of 14mer interaction with N234 and 30mer polyA RNA. CSPs in ^13^C-^1^H HMQC N234 spectra (150uM) for selected residues as a function of 14mer single stranded RNA concentration (red – 0%, orange – 25%, light-blue 100%, dark-blue 200%) as shown in figure 4. Green chemical shift in the presence of 30mer polyA at 1:1 admixture (150uM concentration). Numerous significantly shifted sites lie on a linear trajectory, suggesting that the bound form chemical shift is similar in both cases. Spectra recorded at 850 MHz and 298K.

**Supporting Figure 3.**
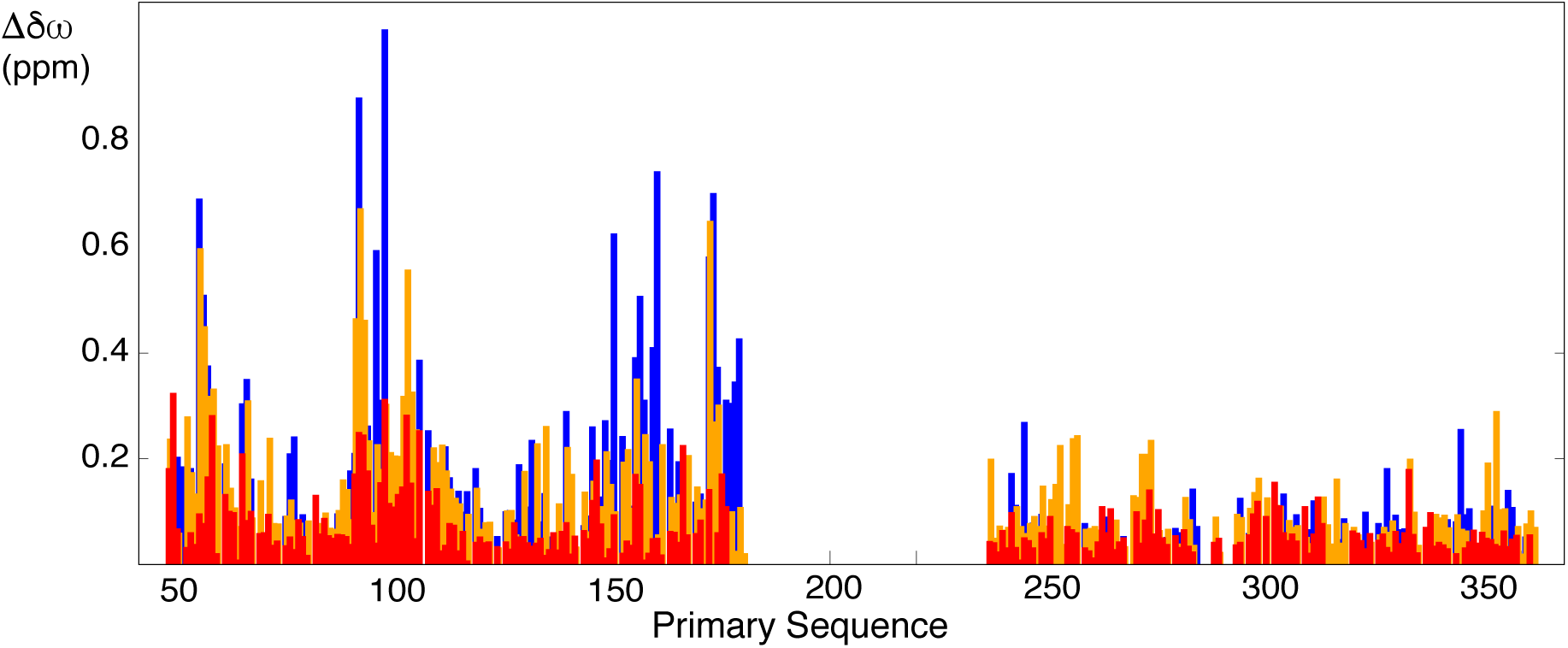
Comparison of ^15^N-^1^H HSQC chemical shifts (compared to non-phosphorylated N234) due to incubation with SRPK1 (red) and GSK-3 (orange) and CK1 (blue) and phosphorylation buffer. Phosphorylated proteins were dialysed into NMR buffer for direct comparison with the non-phosphorylated form.

**Supporting Figure 4.**
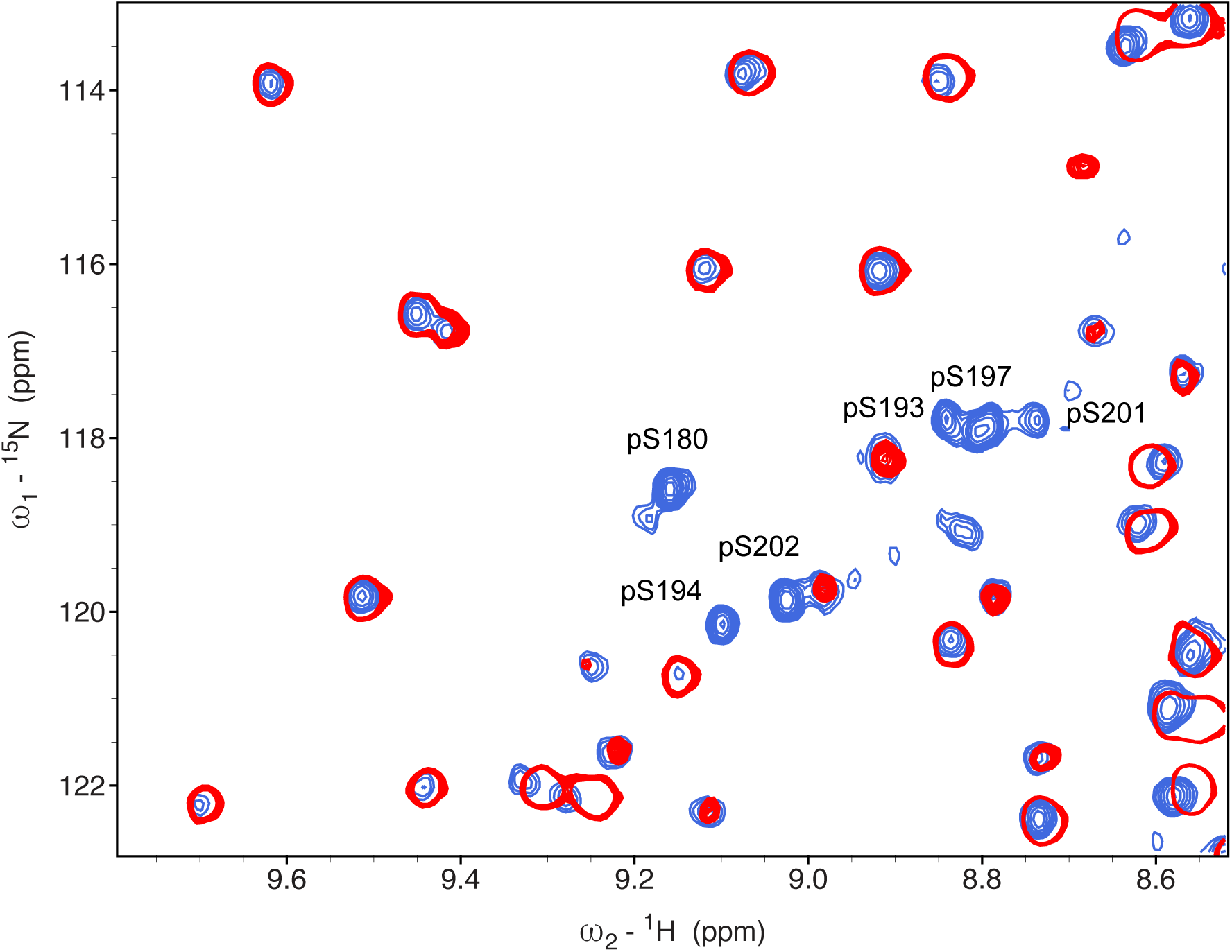
Comparison of ^15^N-^1^H HSQC of unphosphorylated (red) and phosphorylated (blue) N234 following incubation with PKA and phosphorylation buffer. Six resonances appear in the region associated with ^15^N-^1^H cross peaks from phosphorylated residues. These peaks were assigned using standard triple resonance methods.

**Supporting Figure 5.**
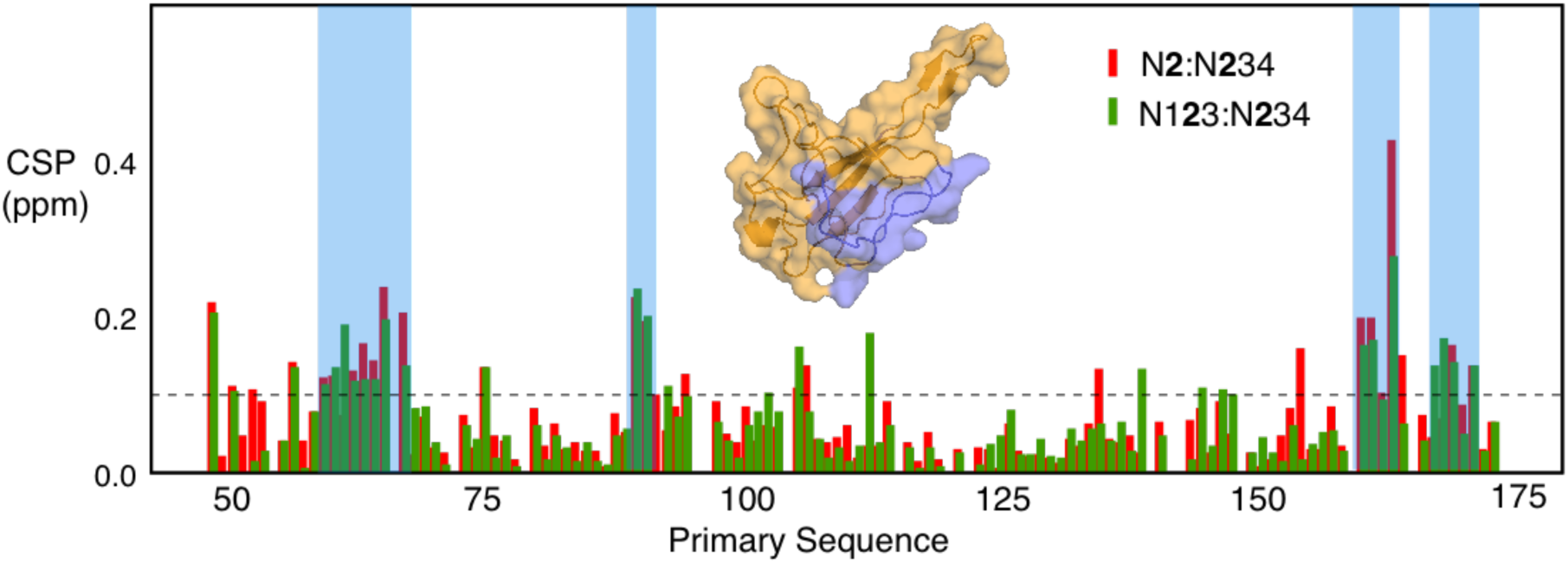
Comparison of ^15^N-^1^H HSQC chemical shifts of N2 in its isolated form and in constructs containing N1 and N3 and N4. Red - difference in chemical shifts of N2 and N2 in N234 Green - difference in chemical shifts of N2 in N234 and N2 in N123 The shading indicates regions where two out of three neighbouring residues show chemical shifts above 0.1ppm. Although small these perturbations are located on a single face of N2 (inset), suggesting that the weak interactions with N4 involve this region.

**Supporting Figure 6.**
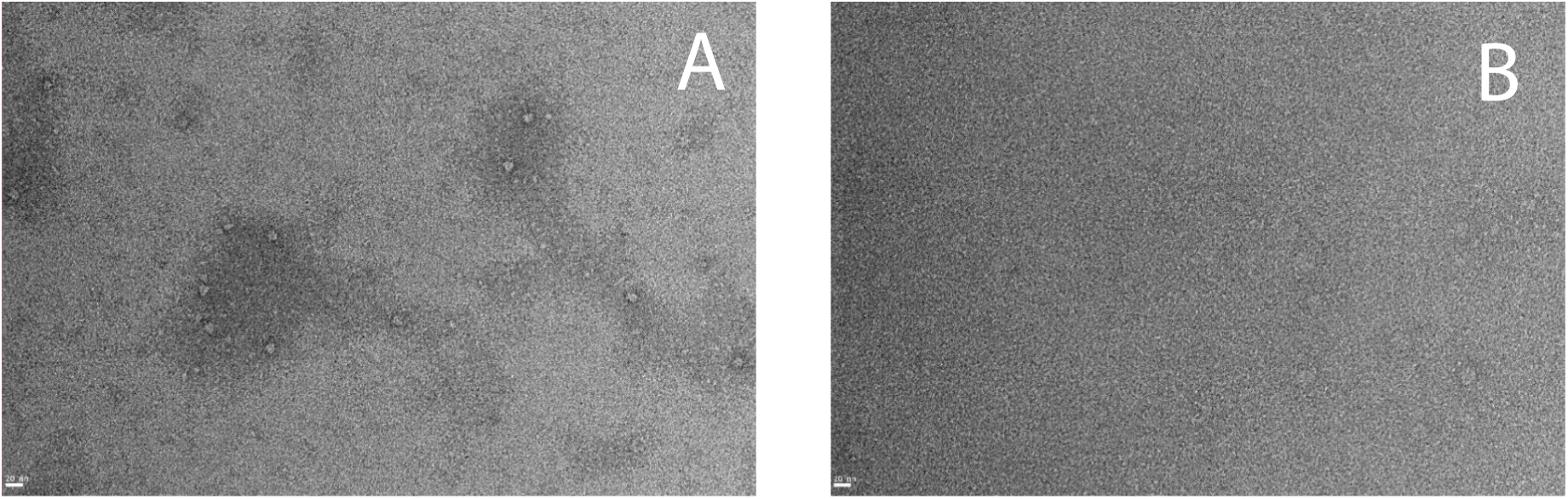
(A) Negative staining electron microscopy of N234 in complex with 14mer RNA showing cage-like particles associated with encapsidation substructures. (B) Such particles are absent in mixtures of RNA with phosphorylated N234.

**Supporting Figure 7.**
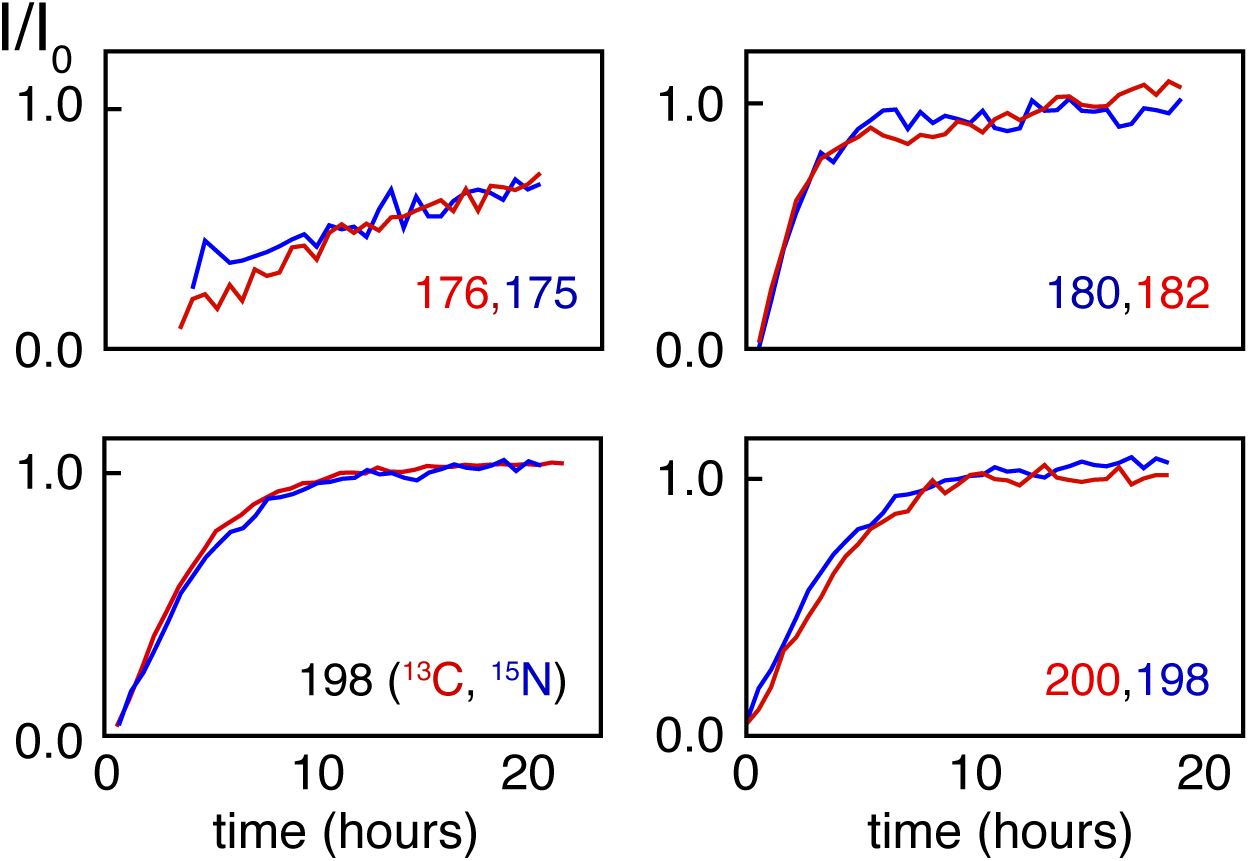
Intensity build-ups for directly phosphorylated amino acids (blue) and neighbouring or near-neighbouring residues (red) are similar. Similarly, the increase in intensity of the sidechain methyl group of T198 mirrors the behaviour of the backbone ^15^N-^1^H correlation peak. These comparisons provide support for the assignment and the kinetics of phosphorylation derived from the analysis of the ^15^N-^1^H correlation peaks of the directly phosphorylated residues.

## Notes

### Competing Interest Statement

The authors have declared no competing interest.

